# De novo whole-genome assembly of *Chrysanthemum makinoi*, a key wild ancestor to hexaploid Chrysanthemum

**DOI:** 10.1101/2021.07.09.451814

**Authors:** N. van Lieshout, M. van Kaauwen, L. Kodde, P. Arens, M.J.M. Smulders, R.G.F. Visser, R. Finkers

## Abstract

Chrysanthemum is among the top ten cut, potted and perennial garden flowers in the world. Despite this, to date, only the genomes of two wild diploid chrysanthemums have been sequenced and assembled. Here we present the most complete and contiguous chrysanthemum *de novo* assembly published so far, as well as a corresponding *ab initio* annotation. The wild diploid *Chrysanthemum makinoi* is thought to be one of the ancestors of the cultivated hexaploid varieties which are currently grown all around the world. Using a combination of Oxford Nanopore long reads, Pacific Biosciences long reads, Illumina short reads, Dovetail sequences and a genetic map, we assembled 3.1 Gb of its sequence into 9 pseudochromosomes, with an N50 of 330 Mb and BUSCO complete score of 92.1%. Our *ab initio* annotation pipeline predicted 95 074 genes and marked 80.0% of the genome as repetitive. This genome assembly of *C. makinoi* provides an important step forward in understanding the chrysanthemum genome, evolution and history.

## INTRODUCTION

As one of the most economically important ornamental crops (Anderson 2007), much time has been invested into understanding *Chrysanthemum x morifolium* and its related varieties and species. One of the key factors of its success as an ornamental crop is the diversity available in petal colors and flower shapes (Song *et al.* 2018), even though the underlying genomic and molecular basis of the shape traits is still poorly understood. This is partly due to the fact that it is a hexaploid with polysomic inheritance (van Geest *et al.* 2017b).

To begin to understand a hexaploid like *C. x morifolium* and its traits, we must first look at the whole genus and research the plant’s origins. The *Chrysanthemum* genus consists of species with a basic number of nine chromosomes but with variable ploidy level, from diploid to decaploid (Wang *et al.* 2014). Native across Eurasia and the northern parts of North America, the genus consists of 40 different species (Liu *et al.* 2012; Liu 2020). More than ten were originally identified as potential source material for the domesticated *C. x morifolium* (Dowrick 1952; Ackerson 1967), including *C. makinoi* (syn. *D. makinoi*), *C. indicum* (syn. *D. indicum*), *C. lavandulifolium* (syn. *D. lavandulifolium*) and *C. zawadskii* (syn. *D. zawadskii*), predominantly in their hexaploid form. The hexaploid *C. vestitum* and tetraploid *C. indicum* were later again suggested as major donors based on comparative morphology, cytology, interspecific hybridization and molecular systematics (Ma *et al.* 2016). Diploids like *C. nankingense*, *C. lavandulifoium* and *C. zawadskii* have also repeatedly been identified as possible contributors (Dai *et al.* 2005; Liu *et al.* 2012; Ma *et al.* 2016). To date, no one has come up with a conclusive model for *C. x morifolium*.

*C. makinoi* is a wild diploid endemic to Japan. While research has been performed in the past with this diploid species (Tanaka 1960; Tanaka and Shimotomai 1968) no one has attempted to assemble its genome. In fact, to date, of the 40 chrysanthemum species only *C. seticuspe* (Hirakawa *et al.* 2019) and *C. nankingense* (Song *et al.* 2018) have whole-genome assemblies. The *C. seticuspe* assembly was made using only short read sequencing and had a total length of 2.722 Gb, with 354 212 contigs, an N50 of 44 741 bps and BUSCO score of 88.8% (Hirakawa *et al.* 2019), while *C. nankingense* was assembled using both long and short reads for a total length of 2.527 Gb, with 24 051 contigs, an N50 of 130 678 bps and BUSCO score of 92.7% (Song *et al.* 2018). Generating a more contiguous assembly of these diploids has been difficult as chrysanthemum genomes are very repetitive and heterozygous (Won *et al.* 2018a; Nguyen *et al.* 2020).

Long read data helps resolve the repetitive sequences and allows for more contiguous contigs to be assembled (van Dijk *et al.* 2018), so we proceeded with an approach that combined both long read, short read, and proximity ligation methods to build a truly robust assembly. This assembly, along with its corresponding organelle assemblies and transcriptome, will expand our understanding of not only the diploid *C. makinoi* but also help illuminate the complicated polyploidization story that led to *C. x morifolium* by providing a robust genomic foundation from which to expand.

## MATERIALS & METHODS

### Plant Material

The *C. makinoi* Matsum. et Nakai or No. JP131333 Ryuunougiku plant, or *C. makinoi* for short, was obtained from the NARO (Tsukuba, Japan) genebank. Cuttings were grown in greenhouses at Wageningen University and Research (WUR-Unifarm) according to standard procedures.

### DNA Extraction, Library Preparation and Sequencing

High molecular weight DNA for long read sequencing was isolated from fresh young *C. makinoi* leaves using a modified (Bernatzky and Tanksley 1986) protocol. Libraries were prepped using the 1D ligation sequencing kits SQK-LSK108 and SQK-LSK109 (Oxford Nanopore Technologies; Oxford, UK) according to the instructions. The samples were sequenced on an Oxford Nanopore GridION using 40 flowcells and the standard protocol. Adaptors were removed using Porechop (Wick 2018) and reads were filtered using Filtlong (Wick 2019), which removed the worst 10% of reads from the shorter reads.

One sample was also sequenced using 4 differently sized insert libraries (270, 350, 400 and 500 bps) and 150 bp paired-end reads on an Illumina Hiseq 2500 (GenomeScan, Leiden, The Netherlands). Samples were processed using the NEBNext® Ultra DNA library Prep Kit from Illumina. Genome characteristics were estimated using Jellyfish v2.2.10 (Marçais and Kingsford 2011) k-mer counts and GenomeScope (Vurture *et al.* 2017).

High molecular weight DNA of *C. makinoi* was also sequenced by GenomeScan across 8 SMRTcells using a PacBio “Sequel SMRT Cell 1M v2” sequencer. Sample preparation was done based on the “PacBio SMRTbell Express Kit v1” protocol. The final library was selected using the Blue Pippin protocol for fragments larger than 15 kb. Primer and polymerase were attached using the “Sequel Binding and Internal Ctrl Kit2.1” kit and purification was done using the PacBio “Procedure & Checklist - AMPure® PB Bead Purication of Polymerase Bound SMRTbell® Complexes” protocol. Sequencing was performed for 10 hours on 7 of the cells and 20 hours for the remaining cell with the recommended amount of “DNA Internal Control Complex 2.1”. The raw data was assessed with the SMRT Link Analysis server v5.1.0.26367 by GenomeScan.

Four tissues (leaves, stems, floral buds, flowers) used in the study were obtained from *a C. makinoi* cultivated in a greenhouse under long-day conditions; 20-hr light/4-hr dark cycle, or under short-day conditions; 11-hr light/13-hr dark cycle, at Dekker Chrysanten (Hensbroek, the Netherlands). All collected plant tissues were frozen immediately in liquid N2 and stored at −70°C until the RNA was extracted and isolated using the RNeasy mini kit (Qiagen; Hilden, Germany) and library prepped using the SQK-PCS109 kit (Oxford Nanopore Technologies; Oxford, UK) according to the instructions. The samples were sequenced separately on an Oxford Nanopore GridION using 9 flowcells in total, according to the standard protocol. Quality control was done using NanoComp v1.9.2 (De Coster *et al.* 2018) and fastq validator from fastq_utils v0.21.0 (Fonseca and Manning) with duplicate read IDs removed.

### Genome Assembly and Scaffolding

Nanopore reads were base-called with Guppy v3.2 (Oxford Nanopore Technologies; Oxford, UK) and filtered to keep only the reads from the “pass” folder (Q≥7) that had a length above 20 Kb and the “fail” folder (Q<7) with a length over 50 Kb. PacBio reads over 30 Kb long were also added into this dataset. This combination of long reads was assembled using SMARTdenovo v1.0.0 (Liu *et al.* 2021) with “generate consensus” set to 1. Purge Haplotigs (Roach *et al.* 2018) was then used to flatten regions of heterozygosity into a single consensus sequence. Illumina data was subsequently used in conjunction with ntEdit v0.9 (Warren *et al.* 2019) in mode 2 and with a K=50 for 2 iterations to polish the contigs. Contiguity was further improved with the use of Hi-C and Chicago proximity ligation methods (Dovetail Genomics; Scotts Valley, USA). Final pseudo-molecule level scaffolding was performed using ALLMAPS v0.9.14 (Tang *et al.* 2015) and an integrated genetic map of hexaploid chrysanthemum (van Geest *et al.* 2017a) (see TABLE S1and FIGURE S1). Some by-hand misassembly corrections, verified with the raw long read data, were also completed (see FIGURE S2). Contigs that remained unplaced among the nine chromosomes in the final assembly were filtered to remove contaminants and unusually high coverage reads. The final chromosomes were named and numbered following the linkage group assignments in a *C. x morifolium* cross found in (van Geest *et al.* 2017a). Read coverage was assessed using Qualimap bamqc v2.2.1 (Okonechnikov *et al.* 2016) and contaminants identified using Centrifuge v1.0.4 (Kim *et al.* 2016) and NCBI’s viral and bacterial libraries (accessed in November 2019). The remaining reads were placed into a chromosome zero with N-gaps of 200 bps in between each contig.

Organelles were assembled by extracting Nanopore and Illumina reads that aligned to the available *C. boreale* chloroplast (Won *et al.* 2018b) and mitochondria (Won *et al.* 2018c) references using Minimap2 v2.17 (Li 2018) and BWA-MEM v0.7.17-r1198-dirty (Li 2013) respectively. A hybrid assembly was then performed for each organelle using Unicycler v0.4.8 (Wick *et al.* 2017). This resulted in a single, circular scaffold assembly for the chloroplast and multiple circular scaffolds for the mitochondria. Based on a visual inspection of each of the mitochondria scaffolds against known chrysanthemum mitochondria assemblies, scaffold 1 was found to represent the entire sequence and was selected as the full circular assembly of the mitochondria genome.

### Genome Analysis and Quality Assessments

QUAST v5.0.2 (Gurevich *et al.* 2013) was used to determine the basic statistics of the final genome assembly such as total length, N50 and the number of contigs/scaffolds. BUSCO v4.0.5 (Simão *et al.* 2015) and the corresponding set of Embryophyta odb10 universal single-copy orthologs was also employed to assess the completeness of the genome.

### Repeat and Transcript Annotation

Before annotating the assembly, we soft-masked the repetitive sequences using RepeatModeler v2.0.1 (Flynn *et al.* 2020).

Gene prediction was done with the Funannotate v1.7.4 (Palmer 2017) pipeline. First the Funannotate pipeline was trained using the cDNA long reads, UniProtKB v2020_04 database (Bateman 2019), and the BUSCO eukaryote odb9 protein database (Simão *et al.* 2015), to create the input dataset for the Funannotate predict pipeline. The predict pipeline was then run with standard settings and the GeneMark-ET, Augustus, GlimmerHMM’, and Snap algorithms. Afterwards filtering of the ab initio gene predictions was done using EVidenceModeler (EVM) (Haas *et al.* 2008).

To functionally annotate the predicted models, an initial comparison was done using blastp v2.6.0 (Camacho *et al.* 2009) against the SWISS-PROT v4 database (Bairoch and Apweiler 2000) with a cut-off e-value of 1.0E-3, word size of 6, max number of hits set to 20 and with the low complexity filter turned on. To identify the domains within the predicted model sets, InterProScan v5.26 (Jones *et al.* 2014) was used along with the panther v12.0 libraries. Finally the results were process by a stand-alone version of Blast2Go (Götz *et al.* 2008) using default settings.

### Data Availability

The final assembly and annotation files for *C. makinoi* Matsum. et Nakai or No. JP131333 Ryuunougiku are available for download at www.chrysanthemumgenome.wur.nl/, along with a genome browser. All the raw data as well as the assembly and annotations files can also be found at ENA under PRJEB44800. The plant accession is available through the NARO genebank.

## RESULTS & DISCUSSION

### Raw Sequence Quality

Nanopore sequencing resulted in 443.25 Gb of data with a read N50 of 22.6 kb. After base calling, removing adaptors and filtering for reads over 20 kb in length from the “pass” folder, which had a Q score >7, and for reads over 50 kb in length from the “fail” folder, the dataset had a coverage of approximately 53x (assuming a haploid genome size of 3.1 GB) and consisted of 3,924,770 reads. Illumina HighSeq yielded 113.2, 142.0, 133.7 and 120.0 Gb of raw data for the 270, 350, 400 and 500 bp insert size libraries respectively. Between 90.5-94.6% of reads in each insert size had a quality “q” score greater or equal to 30. PacBio sequencing resulted in 70 Gb of data with an average subread length of 15.5 kb and a N50 of 24.1 kb. This meant a coverage of approximately 30.6x (assuming a haploid genome size of 3.1 GB).

The nanopore cDNA sequencing resulted in datasets with 4.8 – 7.9 million reads, an average N50 of 1.2-1.4 kb and between 5.0-7.9 Gb total (see TABLE 1).

**TABLE 1.** SEQUENCING DETAILS OF CDNA FROM DIFFERENT PLANT ORGANS IN C. MAKINOI.

| Source | Mean read length (b): | Mean read quality: | Number of reads: | Read length N50 (b): | Total bases: |
| --- | --- | --- | --- | --- | --- |
| Leaf (short day) | 977.0 | 9.0 | 7,587,930 | 1,189 | 7,413,732,394 |
| Leaf (long day) | 973.0 | 9.4 | 6,780,899 | 1,209 | 6,597,664,366 |
| Calyx | 1,040.2 | 10.0 | 4,833,397 | 1,247 | 5,027,548,849 |
| Flower Disk<br>Florets | 1,012.1 | 9.5 | 7,072,131 | 1,331 | 7,157,901,095 |
| Flower Buds | 993.6 | 9.1 | 7,917,800 | 1,256 | 7,867,489,497 |
| Flower Ray<br>Florets | 1,002.8 | 9.6 | 7,000,372 | 1,250 | 7,020,311,566 |
| Meristem | 1,048.0 | 10.2 | 5,075,164 | 1,263 | 5,318,808,662 |
| Stem (short day) | 997.1 | 9.5 | 7,936,023 | 1,232 | 7,912,613,286 |
| Root | 1,060 | 8.4 | 5,272,384 | 1,389 | 5,591,241,404 |

### Genome Size and Characteristics

K-mers (K=31) were extracted from the paired-end HiSeq Illumina reads, counted using Jellyfish v2.2.10 (Marçais and Kingsford 2011) and analyzed with GenomeScope (Vurture *et al.* 2017) to estimate the genome haploid length, heterozygosity and repeat content. The analysis converged and estimated a haploid genome size of 1.72 Gb, a heterozygosity of 1.51% (this value ranges from ~0-2% (Vurture *et al.* 2017)) and marked 53.6% of the genome as unique (Figure 1). This indicates that the genome is repetitive and highly heterozygous. The haploid genome size of the chrysanthemum diploids has been estimated between 2.90 ± 0.03 Gb for *C. seticuspe* (Hirakawa *et al.* 2019) and 3.24 Gb for *C. nankingense* (Song *et al.* 2018) using flow-cytometry. The Genome Size Asteraceae Database (GSAD) estimates an average 1C of 3.82 GB for chrysanthemum using flow-cytometry, though this is likely an overestimation as the median is 3.1 GB (Garnatje *et al.* 2011). It is known that sequence based genome estimation methods underestimate genome size (Pflug *et al.* 2020) with GenomeScope being particularly sensitive to the k-mer count cut-off parameter (Vurture *et al.* 2017). This parameter is meant to distinguish repetitive sequences from organelle sequences, so that the repetitive k-mers are used to calculate the genome size while organelle k-mers are discarded, but this becomes impossible if the repetitive sequence k-mers are as abundant as the organelle k-mers (Vurture *et al.* 2017). With the high level of heterozygosity indicated by GenomeScope and confirmed with later analyses, it would be difficult to distinguish these k-mers from each other, resulting in many of the repetitive region k-mers also being discarded and producing a substantially underestimated genome size. We expect a true genome size closer to the cytometry predictions of *C. nankingense* and *C. seticuspe*.

**FIGURE 1.**
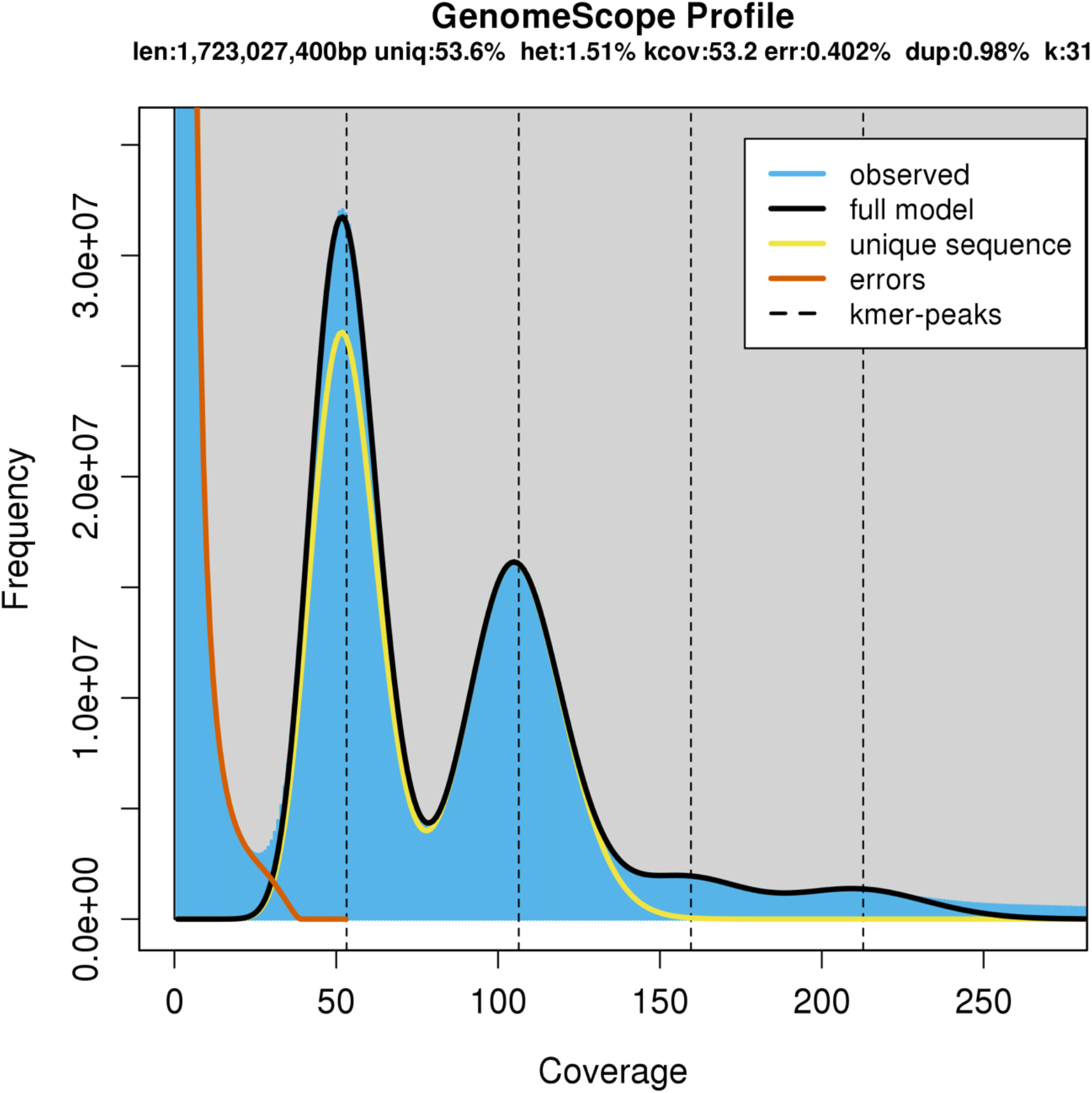
A K-MER (K=31) DISTRIBUTION BASED ON THE ILLUMINA READS, MODELED AND VISUALIZED USING GENOMESCOPE.

### Genome Assembly and Quality

After initial assembly with SMARTdenovo (Ruan *et al.*) we had 39,105 contigs, spanning 4.1 Gb and with an N50 of 139.2 kb. Purge Haplotigs (Roach *et al.* 2018) produced a flattened assembly of 15 236 contigs, spanning 3.1 Gb, with an N50 of 255.8 kb. After two rounds of polishing with ntEdit v0.9 (Warren *et al.* 2019) using Illumina data, the assembly size was 3.1 Gb, and made up of 15,226 contigs and with an N50 of 258.2 kb.

To scaffold the contigs, maps were generated using Hi-C and Chicago proximity ligation methods (Dovetail Genomics; Scotts Valley, USA). This method generated 4 254 scaffolds, covering a total length of 3.1 Gb and with an N50 of 168.9 Mb. The assembly was further superscaffolded into pseudochromosomes using ALLMAPS v0.9.14 (Tang *et al.* 2015) using a genetic map from a hexaploid *C. moriflorium* (van Geest *et al.* 2017a). This resulted in a final assembly that was 3.1 Gb long, scaffolded into 9 pseudochromosomes and with an N50 and L50 of 330.0 Mb and 5 scaffolds respectively (Table 2.).

**TABLE 2.**
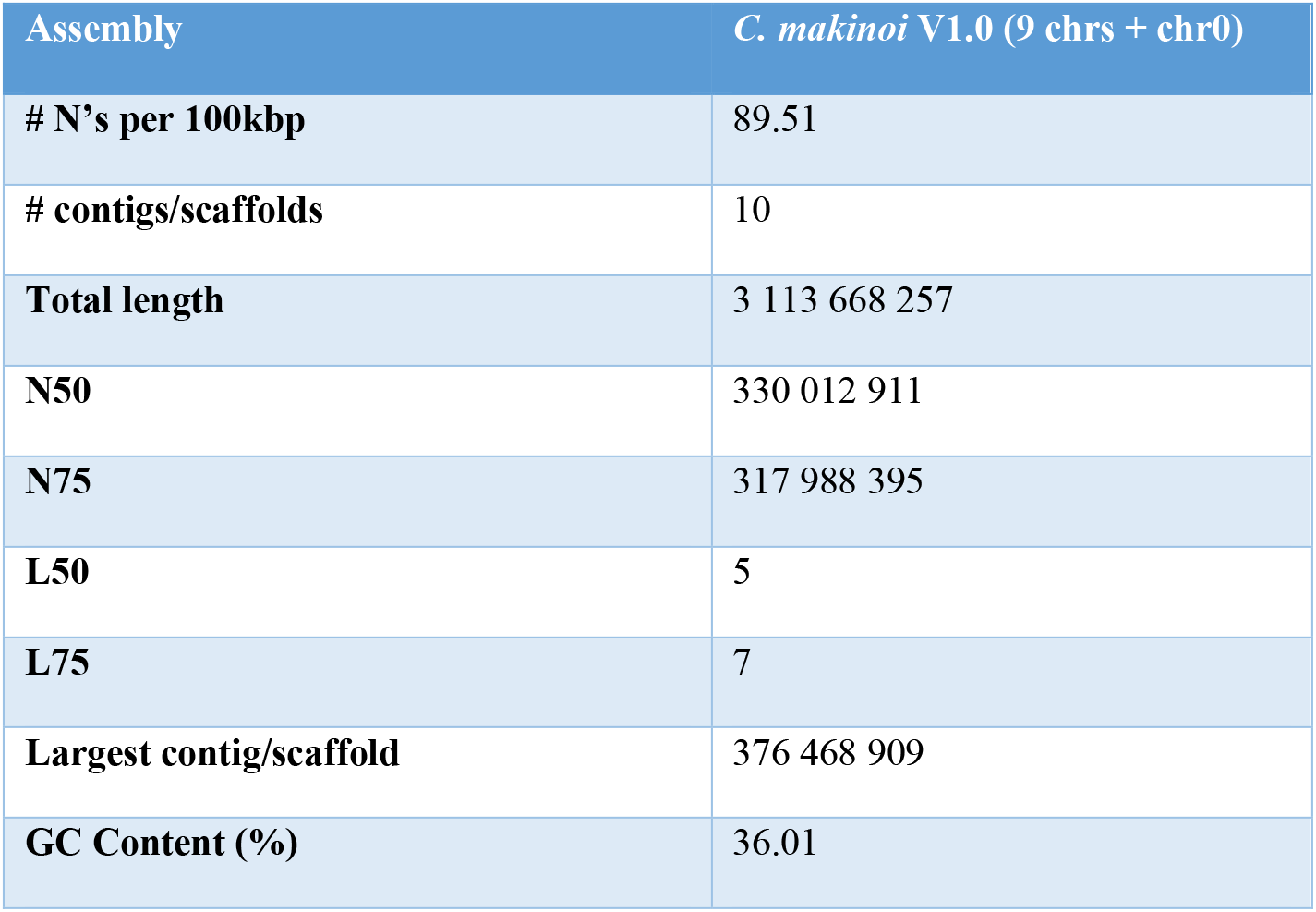
C. MAKINOI DE NOVO GENOME ASSEMBLY METRICS ESTIMATED USING QUAST.

| Assembly | *C. makinoi* V1.0 (9 chrs + chr0) |
| --- | --- |
| # N's per 100kbp | 89.51 |
| # contigs/scaffolds | 10 |
| Total length | 3 113 668 257 |
| N50 | 330 012 911 |
| N75 | 317 988 395 |
| L50 | 5 |
| L75 | 7 |
| Largest contig/scaffold | 376 468 909 |
| GC Content (%) | 36.01 |

The unplaced contigs were curated before being placed into chromosome 0 using the classification engine Centrifuge v1.0.4 (Kim *et al.* 2016). Of the 4 206 unplaced contigs, 824 were marked as coming from a non-eukaryote source and removed. The Illumina reads were also aligned back to all the contigs using Minimap2 v2.17 (Li 2018) and then their coverage was assessed using Qualimap v2.2.1 (Okonechnikov *et al.* 2016). Contigs with a coverage higher than one standard deviation from the average were removed. This resulted in a final set of 3 337 contigs, covering a total of 198.3 Mb, which were placed into chromosome 0.

BUSCO scores, which provide a set of universal single-copy orthologs, were also used to assess the completeness of the assemblies (see Table 3). Using the Embryophyta odb10 set with BUSCO v4.0.5 (Simão *et al.* 2015), the final assembly had a complete BUSCO score of 92.1% indicating a high overall quality. A full breakdown of the BUSCO score can be seen in Table 3.

**TABLE 3.**
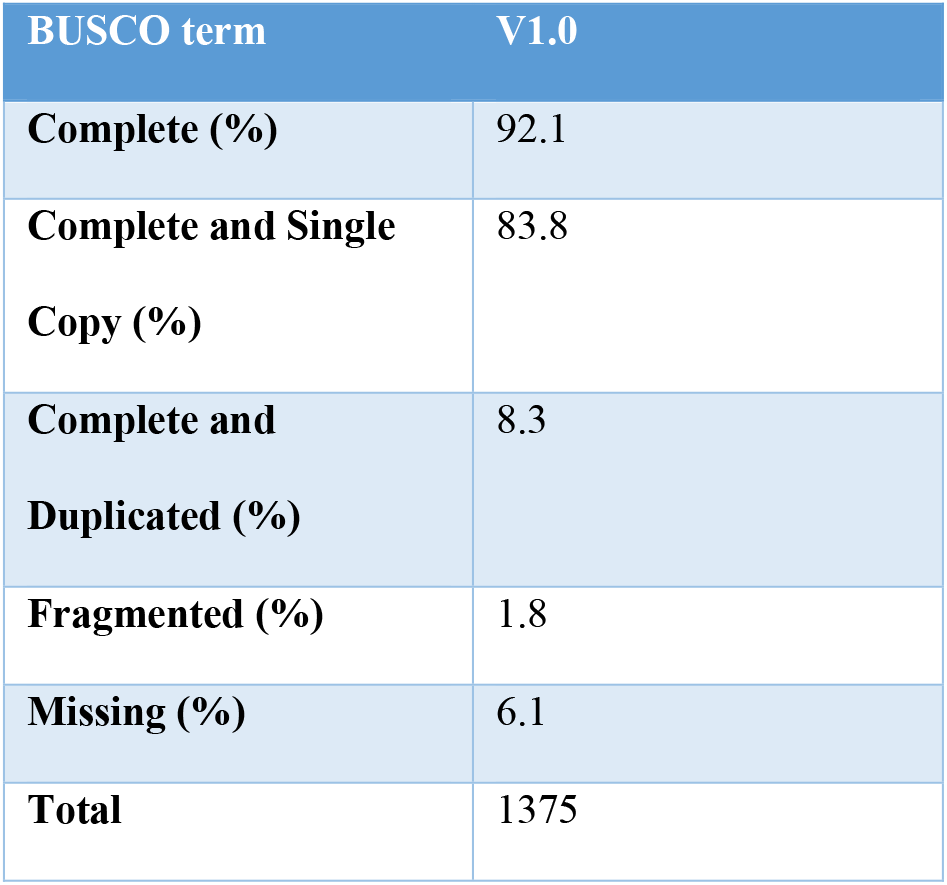
OUTPUT FROM THE BUSCO ANALYSIS PIPELINE TO ASSESS GENE COMPLEMENT COMPLETENESS.

| BUSCO term | V1.0 |
| --- | --- |
| Complete (%) | 92.1 |
| Complete and Single Copy (%) | 83.8 |
| Complete and Duplicated (%) | 8.3 |
| Fragmented (%) | 1.8 |
| Missing (%) | 6.1 |
| Total | 1375 |

For comparison, the exclusively short read based assembly of *C. seticuspe* had a total length of 2.722 Gb, with 354 212 contigs, an N50 of 44.7 kb and BUSCO score of 88.8% (Hirakawa *et al.* 2019). The *C. nankingense* assembly had a total length of 2.527 Gb, with 24 051 contigs, an N50 of 130.7 kb and BUSCO score of 92.7% (Song *et al.* 2018). Thus we were able to produce a substantially more contiguous assembly without sacrificing completeness.

### Repetitive Regions

Using RepeatModeler (Flynn *et al.* 2020), 80.04% of the genome was marked as repetitive. Large genomes have accumulated repeats (Kelly and Leitch 2011) and the k-mer analysis already indicated we were dealing with a largely repetitive genome. Of the 6 799 identified repeat families in *C. makinoi*, 76.6% were identified as long terminal repeats (LTRs). Of the LTRs, 27.1% could be identified as *Copia* and 7.4% as *Gypsy*. A similar analysis in *C. nankingense* marked 69.6% of their assembly as repetitive and found LTRs to make up 67.7% of the identified tandem repeats, with 36.5% being *Copia* and 30.9% being *Gypsy* (Song *et al.* 2018). The lower rate of repetitiveness and identified LTRs in *C. nankingense* may be due to the difference in contiguity, with *C. nankingense* consisting of over 24 000 contigs (Song *et al.* 2018) to our 9 pseudochromosomes and 3 337 unplaced contigs, as it has been shown that more complete genome assemblies will identify more LTRs (Ou *et al.* 2018). Analysis of various Asteraceae has shown fluctuations between members in relative abundance of *Copia* vs *Gypsy*, with sunflower (*Helianthus annuus*) amplifying *Gyspy* over *Copia* (Cavallini *et al.* 2010; Buti *et al.* 2011; Natali *et al.* 2013; Giordani *et al.* 2014; Badouin *et al.* 2017) while horseweed *(Conyza canadensis*) and globe artichoke (*Cynara Cardunculus* var. scolymus) showed the reverse (Peng *et al.* 2014; Scaglione *et al.* 2016). Earlier studies with *C. nankingense* and *C. boreale* suggested that in chrysanthemum the abundances of *Copia* and *Gypsy* were similar, with *Copia* being slightly more abundant and undergoing amplification slightly earlier (Won *et al.* 2018a; Song *et al.* 2018) but our results suggest that, at least in *C. makinoi,* there is a more substantial difference in abundance, like that seen in horseweed and globe artichoke. A systematic analysis of a variety of chrysanthemum species at various ploidy levels should be undertaken to gain better insight as these repeat types are a known driving force of plant genome evolution (Todorovska 2007).

### Transcript Annotation

Each algorithm in the Funannotate (Palmer 2017) pipeline produced a set of ab initio gene models (see TABLE S2). The evidence for each gene model was weighed using an EVM approach and identified 95 064 ab initio predicted gene models. This is higher than the plant average of 36 795 (Ramírez-Sánchez *et al.* 2016) but could be explained by the presence of pseudogenes (Xiao *et al.* 2016). Other Asteraceae including *Artemisia annua* (63 226 gene models) (Shen *et al.* 2018), sunflower (52 232 gene models) (Badouin *et al.* 2017), *Mikania micrantha* (46 351 gene models) (Liu *et al.* 2020), and *C. seticuspe* (71 057 gene models) (Hirakawa *et al.* 2019), and *C. nankingense* (56 870 gene models) (Song *et al.* 2018) also have substantially more than the average number of gene models. To investigate this further, an analysis of the structure and length of the annotated genes was also performed. The genes had an average coding sequence length of 876 bps and a maximum of 12 735 bps. This is shorter than the average plant gene length of 1 308 bps but within the first quartile of average plant gene length (Ramírez-Sánchez *et al.* 2016). In line with the finding that plants tend to have less exons per protein than other organisms (Ramírez-Sánchez *et al.* 2016), 15.9% of the genes in *C. makinoi* were found to consist of a single exon. The average intron length within our gene set was found to be 446 bps, with a range of 11-19 668 bps and median of 140 bps. This indicates that the majority of introns are relatively small. The distribution is similar to what has been found in maize (which had a mean of 516 bps and median of 146 bps) (Schnable *et al.* 2009).

Typically transposable elements accumulate in the centromeric and pericentromeric regions as they establish, maintain and stabilize the centromeres of eukaryotes (Klein and O’Neill 2018). Thus one can estimate the centromeric region of a chromosome based on a low gene density (Figure 2.; red ring) and high repetitive sequence density (Figure 2.; orange ring) but this pattern is not visible in *C. makinoi* as both the genes (red ring) and repetitive sequences (orange ring) are evenly distributed across. In fact, instead of clustering by region, the repetitive sequence density in *C. makinoi* has a positive Pearson Correlation value of 0.60 with gene density. A possible explanation for this correlation is that chrysanthemum, like other Asteraceae, have LTRs driving a lot of diversity (Wang *et al.* 2014). Each LTR family has it’s own distribution characteristics in plant genomes (Chen 2007; Zhanga *et al.* 2014) and LTRs make up 76.6% of the identified repeat families in *C. makinoi*. The sheer volume of the LTRs that distribute in gene rich areas could be overwhelming the signal of repetitive sequences with a centromeric/pericentromeric preference. This is further supported by previous work on repetitive elements in *C. boreale* which found, using optical techniques, a strong enrichment for LTRs and that the majority of repetitive sequences identified did not show a preference for centromeric or peri-centromeric regions (Won *et al.* 2018a).

**FIGURE 2.**
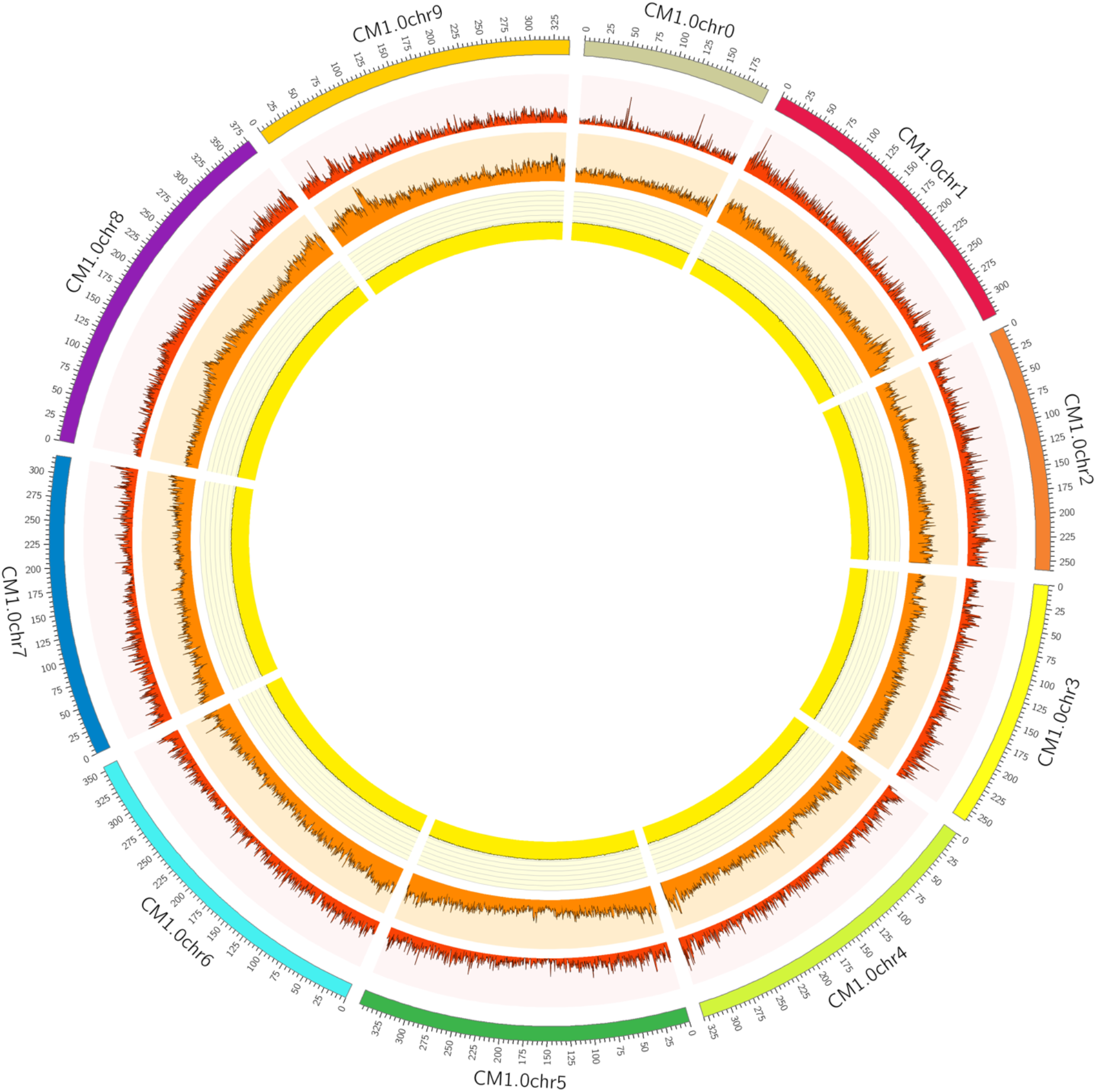
CIRCOS PLOT SHOWING THE PSEUDOMOLECULES (OUTER RING), GENE DENSITY (RED RING), REPETITIVE ELEMENT CONTENT (ORANGE RING) AND GC CONTENT (YELLOW RING) WITH A BIN SIZE OF 500KB.

Blast2Go (Götz *et al.* 2008) was used to functionally annotate the final gene model set. From our predicted gene models, 11.0% were assigned a putative functional label and 2.9% an enzyme code. Looking at the GO-level distribution, the majority of the gene models that were annotated as relating to a biological process (P) or molecular function (F) could not be identified to a high level of specificity, except the cellular component (C) annotated genes (see FIGURE S3). This means that Blast2Go struggled to be more specific about the function of the identified biological process genes beyond i.e. “nitrogen compound metabolic process” but could get much more specific with the cellular component annotated genes.

## CONCLUSION

Having assembled the most complete and contiguous chrysanthemum genome available to date we have made an important step forward in our understanding of the genomics of this complex important ornamental crop. This reference will provide a guide for further research in chrysanthemum breeding traits, origin and strategies for assembling related higher ploidy varieties. This genome can act as a reference to assist in the ordering of other diploid chrysanthemum sequences as well as help reduce the complexity of assembly in closely related polyploids as has been done in several other species (Lukaszewski *et al.* 2014; Li *et al.* 2015; Bertioli *et al.* 2016; Kyriakidou *et al.* 2018; Edger *et al.* 2019).

## ACKNOWLEDGEMENTS

We would like to acknowledge and thank our industry partners, Deliflor Chrysanten, Dekker Chrysanten, Royal Van Zanten and Dümmen Orange as well as TKI T&U for their funding and support under TKI project KV1605-114. The National Agriculture and Food Organization (Japan) is also kindly acknowledged for sharing the *Chrysanthemum makinoi* plant accession JP131333 with us for this project.

## SUPPLEMENTARY TABLES AND FIGURES

**TABLE S1.** ALLMAPS SUMMARY BETWEEN THE HEXAPLOID INTEGRATED GENETIC MAP AND C. MAKINOI SCAFFOLDS THAT DETAILS THE MARKER STATS FOR THE SEQUENCES THAT WERE ANCHORED, ONLY ORIENTED OR UNPLACED.

|  | Anchored | Oriented | Unplaced |
| --- | --- | --- | --- |
| Markers (unique) | 48 482 | 48 393 | 910 |
| Markers per Mb | 16.6 | 16.6 | 4.0 |
| N50 Scaffolds | 8 | 8 | 0 |
| Scaffolds | 81 | 63 | 4 238 |
| Scaffolds with 1 marker | 8 | 0 | 105 |
| Scaffolds with 2 markers | 7 | 5 | 62 |
| Scaffolds with 3 markers | 2 | 2 | 28 |
| Scaffolds with ≥4 markers | 64 | 56 | 72 |
| Total bases | 2 914 525 386 (92.7%) | 2 912 184 103 (92.6%) | 229 073 915 (7.3%) |

**TABLE S2.** NUMBER OF GENE MODELS PRODUCED BY EACH ALGORITHM AS PART OF THE FUNANNOTATE PIPELINE.

| Algorithm | Number of Models |
| --- | --- |
| Genemark | 212 885 |
| Augustus | 81 180 |
| High Quality Augustus | 23 894 |
| GlimmerHMM' | 977 715 |
| Snap | 557 986 |

**FIGURE S1.**
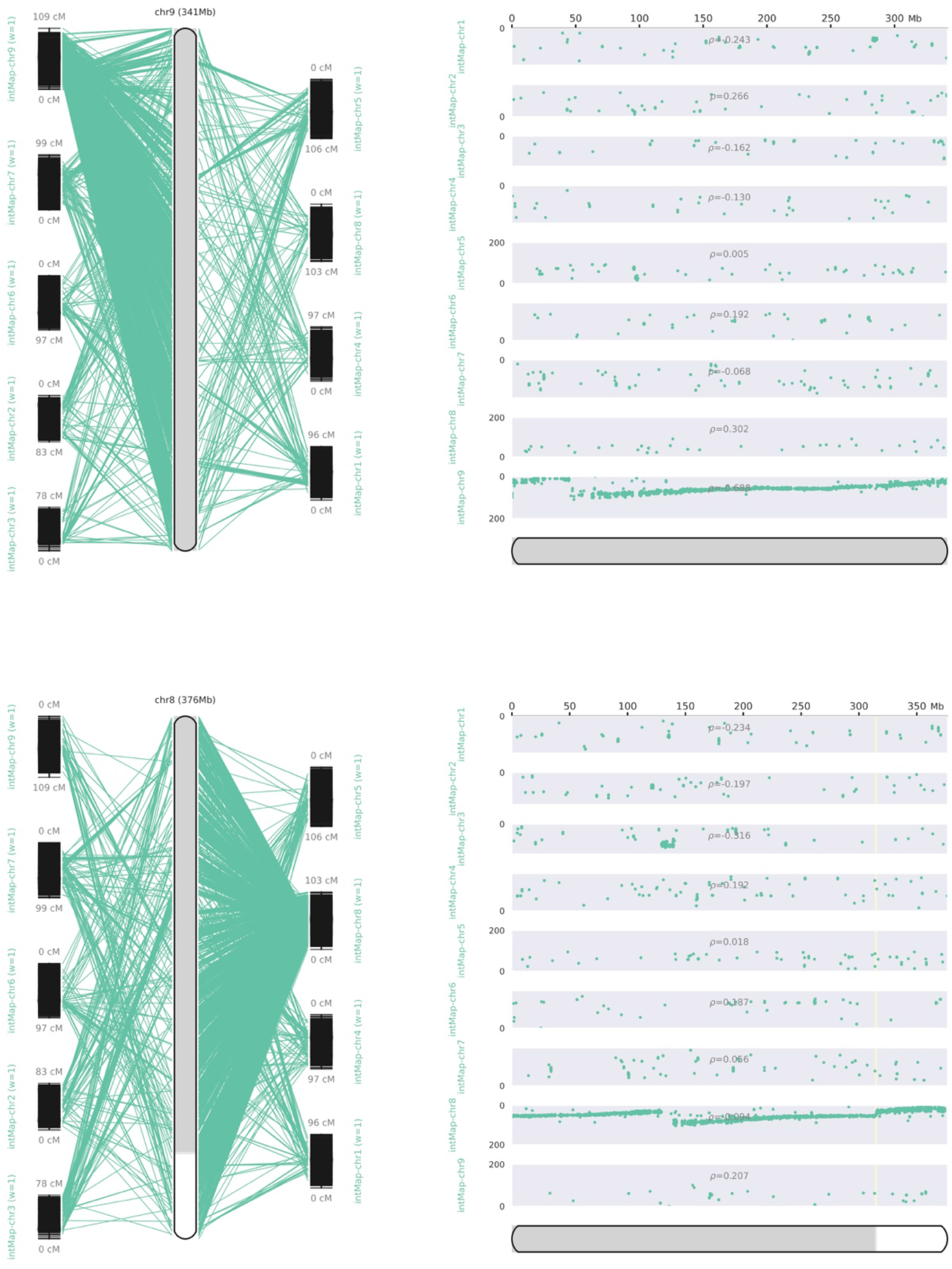
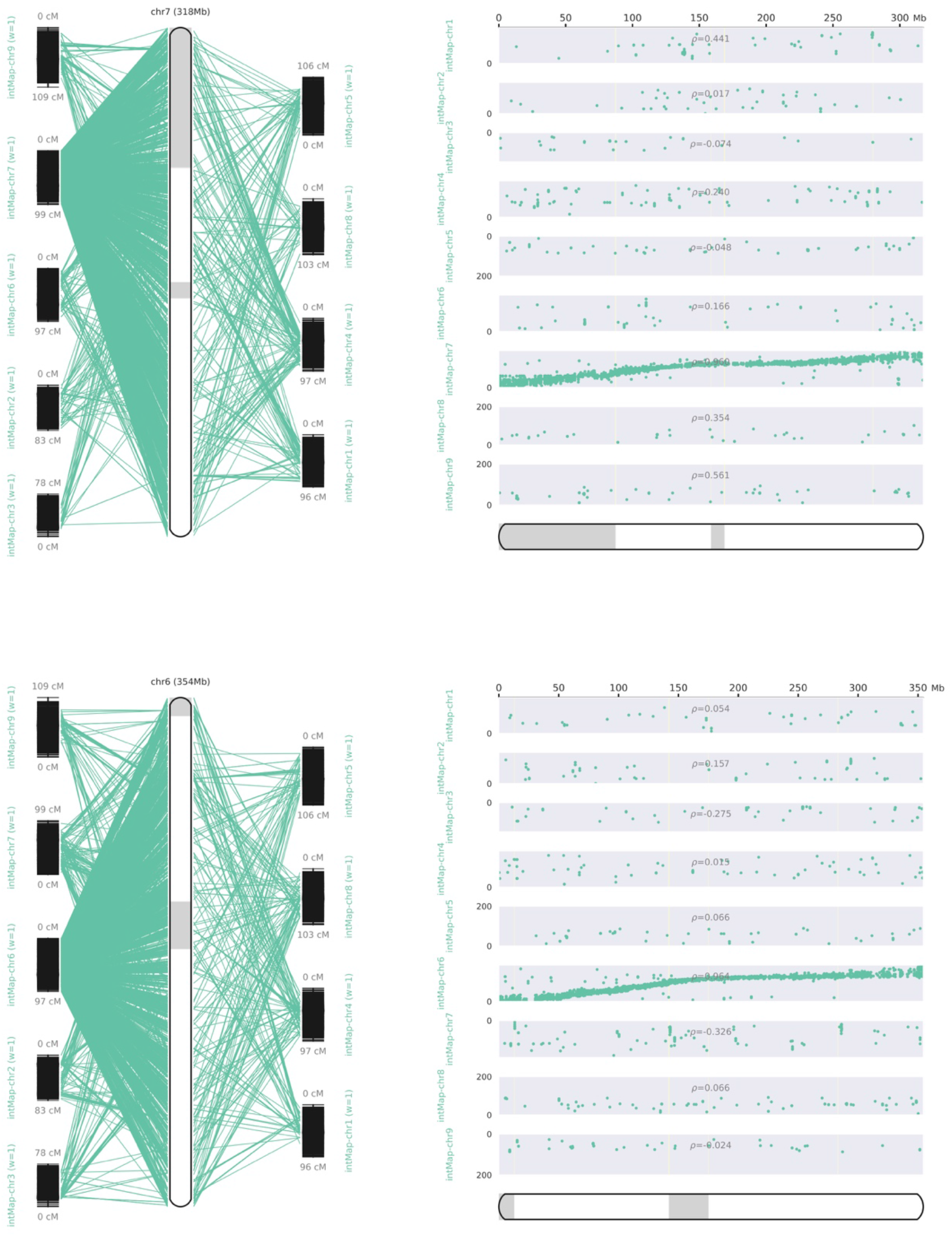
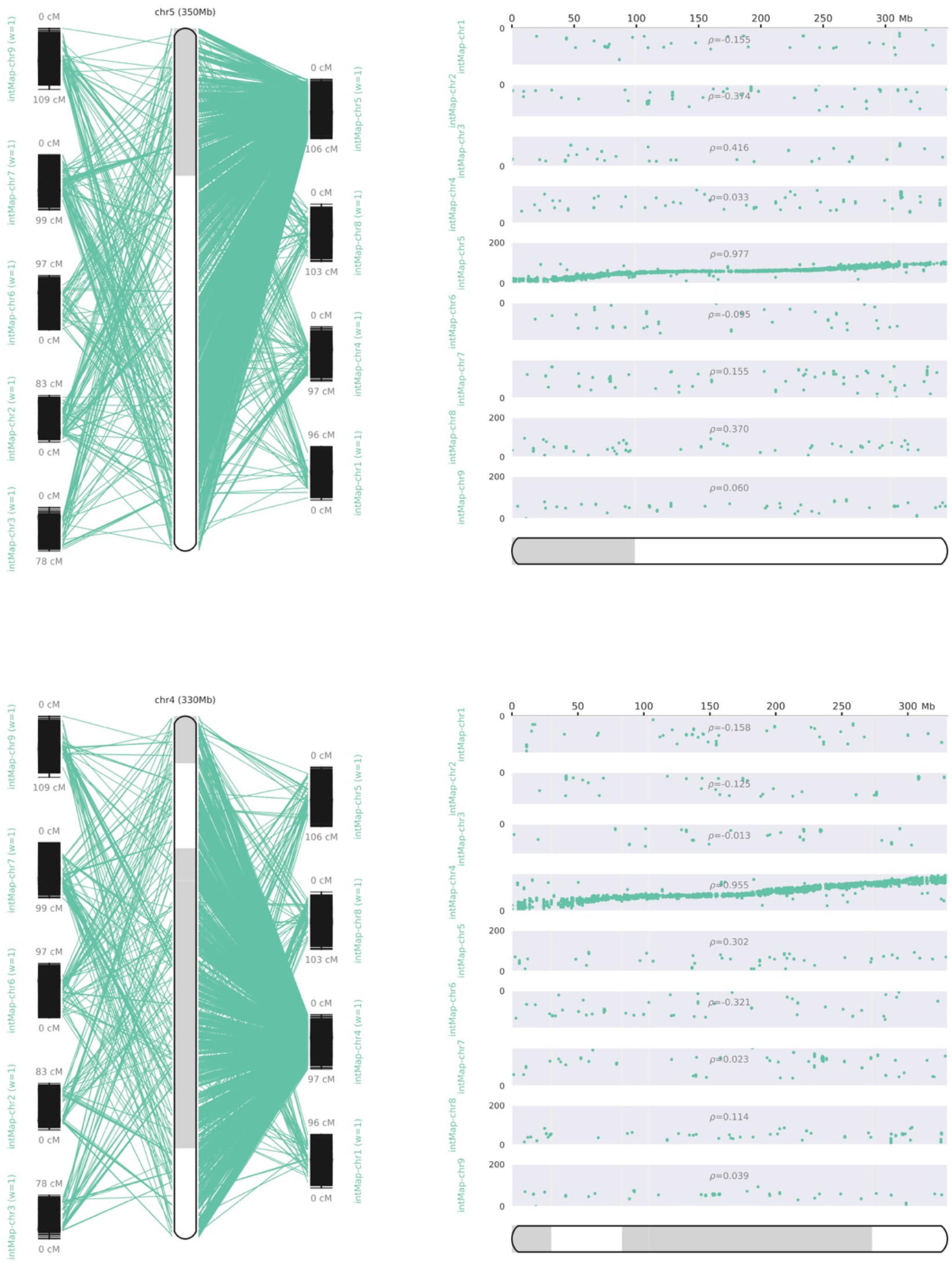
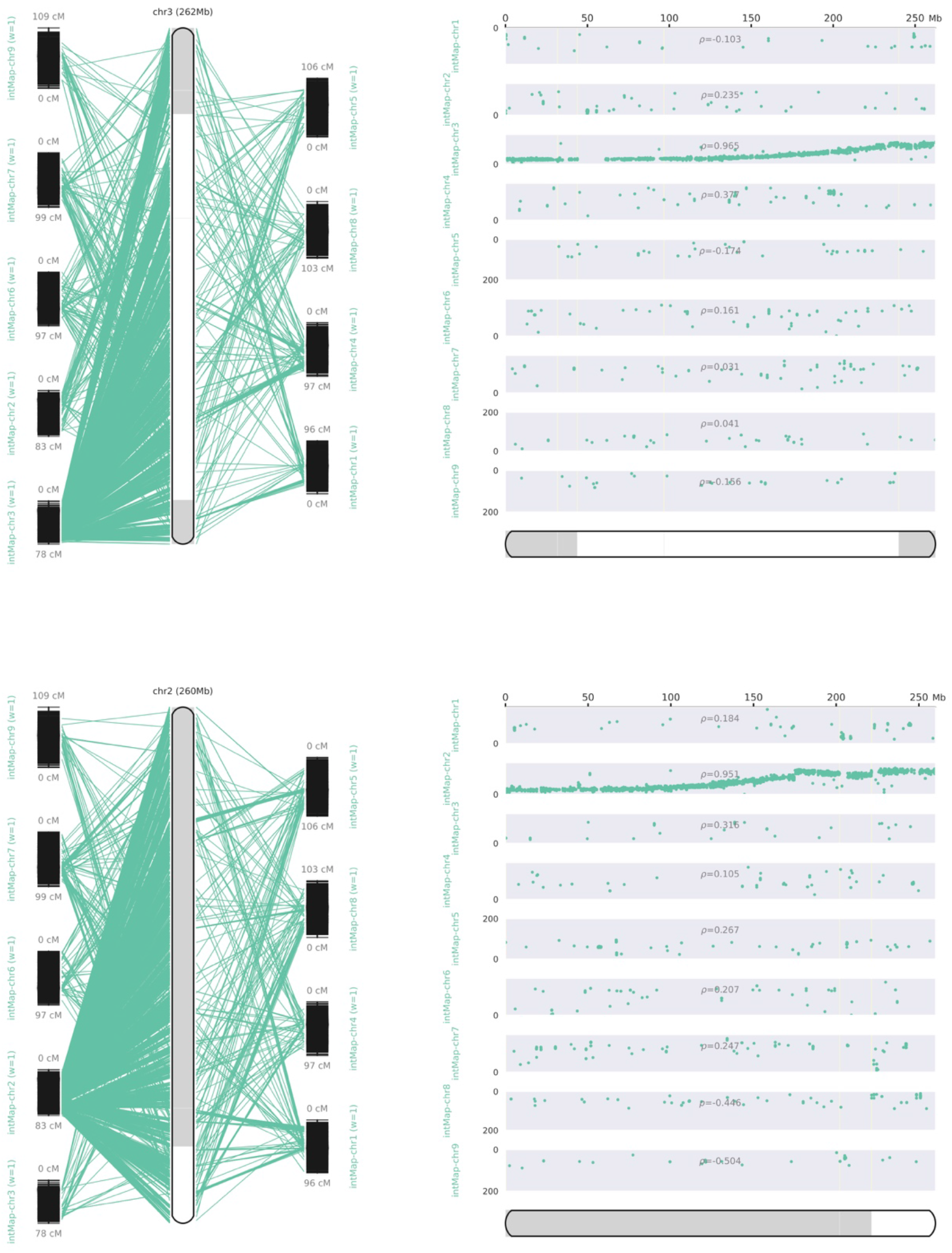
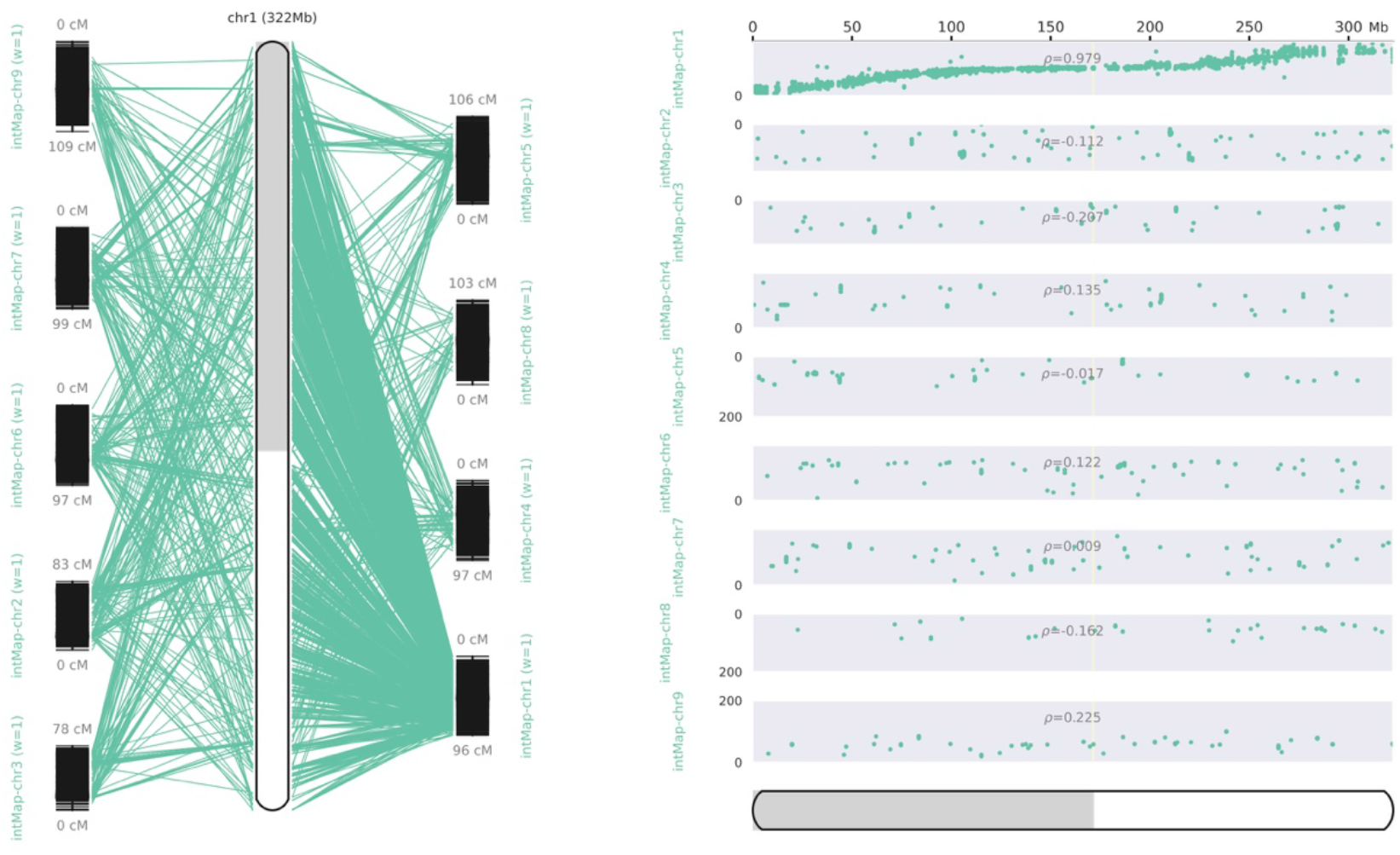
ALLMAPS CHROMOSOME RECONSTRUCTION PLOTS OF C. MAKINOI USING A HEXAPLOID INTEGRATED GENETIC MAP, WHERE THE LEFT PLOT CONTAINS A SIDE BY SIDE ALIGNMENT OF CHROMOSOME AND LINKAGE GROUPS AND THE RIGHT SCATTERPLOT, THE PHYSICAL VS MAP LOCATIONS OF THE MARKERS. THE PHYSICAL LOCATIONS OF THE MARKERS WERE DETERMINED USING BLAST, USING THE BEST HIT. A SMALL SUBSET OF THE OF MARKERS WERE INCORRECTLY PLACED BUT THE HIGH DENSITY OF THE MAP PROVIDED AMPLE COVERAGE FOR ALLMAPS TO COME TO THE CORRECT CONSENSUS ANCHORING.

**FIGURE S2.**
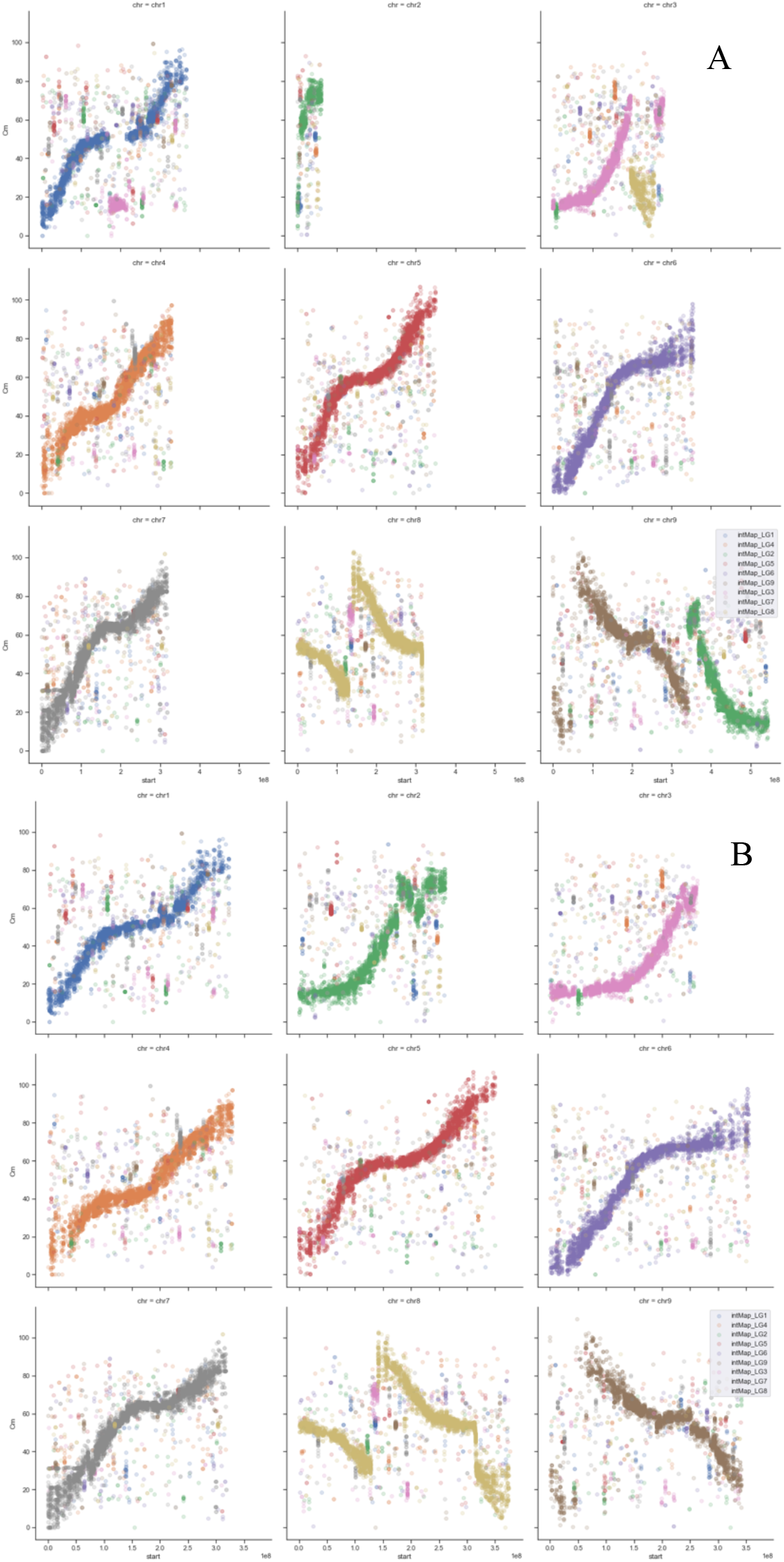
ALLMAP VISUALIZATION OF PSEUDOCHROMOSOME ASSEMBLIES AGAINST A HEXAPLOID GENETIC MAP BEFORE (A) AND AFTER (B) ONT VERIFIED BY-HAND CORRECTIONS.

**FIGURE S3.**
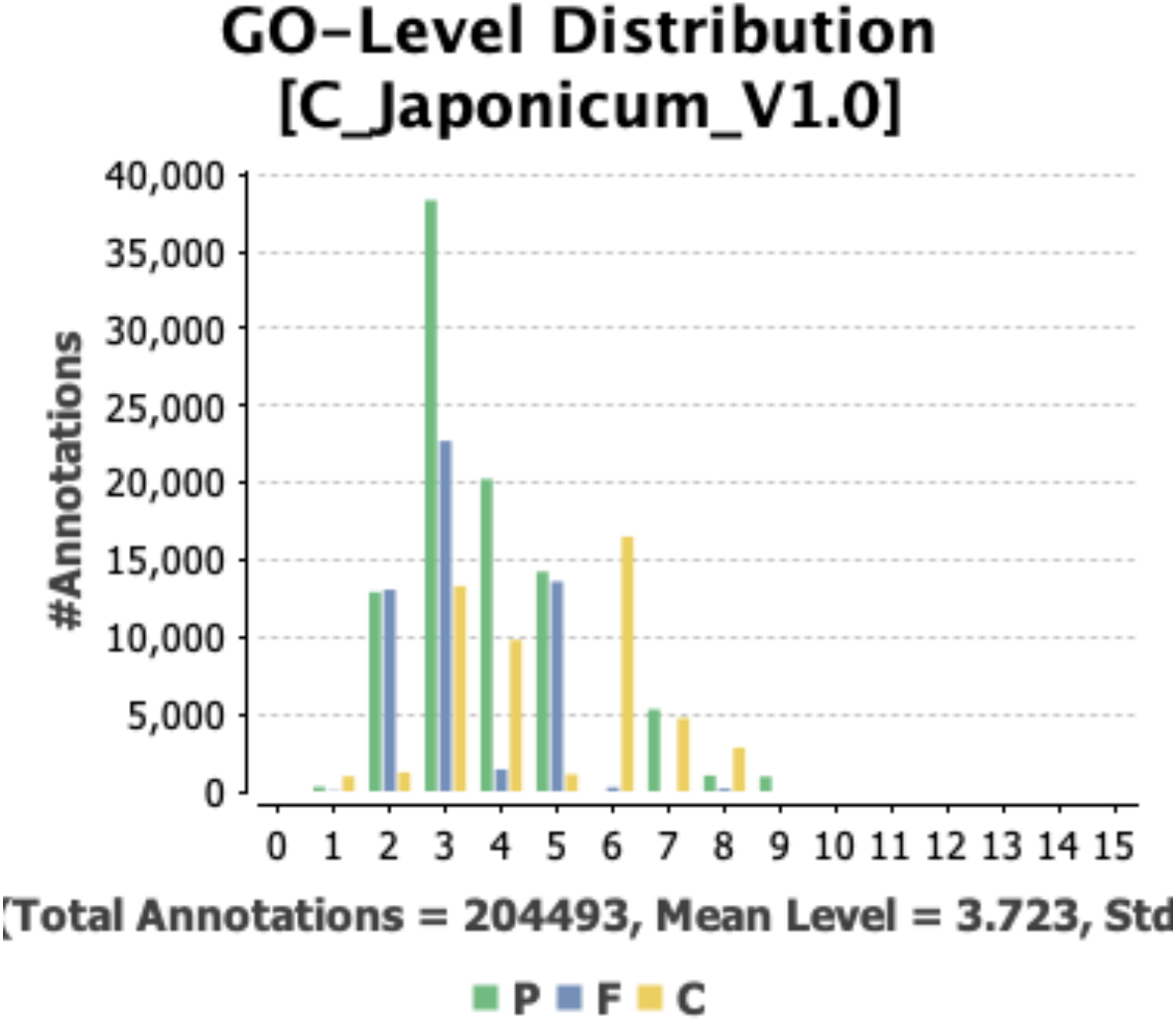
GO-LEVEL DISTRIBUTIONS OF SPECIFICITY FROM THE C. MAKINOI ANNOTATIONS SORTED BY FUNCTION.

## LITERATURE CITED

Ackerson, C., 1967 Original species of the chrysanthemum. Natl. Chrysanth. Soc. Bull. 23: 105–107.

Anderson, N. O., 2007 Chrysanthemum, pp. 389–437 in Flower Breeding and Genetics, Springer Netherlands.

Badouin, H., J. Gouzy, C. J. Grassa, F. Murat, S. E. Staton et al., 2017 The sunflower genome provides insights into oil metabolism, flowering and Asterid evolution. Nature 546: 148–152.

Bairoch, A., and R. Apweiler, 2000 The SWISS-PROT protein sequence database and its supplement TrEMBL in 2000. Nucleic Acids Res. 28: 45–48.

Bateman, A., 2019 UniProt: A worldwide hub of protein knowledge. Nucleic Acids Res. 47: D506–D515.

Bernatzky, R., and S. D. Tanksley, 1986 Genetics of actin-related sequences in tomato. Theor. Appl. Genet. 72: 314–321.

Bertioli, D. J., S. B. Cannon, L. Froenicke, G. Huang, A. D. Farmer et al., 2016 The genome sequences of Arachis duranensis and Arachis ipaensis, the diploid ancestors of cultivated peanut. Nat. Genet. 48: 438–446.

Buti, M., T. Giordani, F. Cattonaro, R. M. Cossu, L. Pistelli et al., 2011 Temporal dynamics in the evolution of the sunflower genome as revealed by sequencing and annotation of three large genomic regions. Theor. Appl. Genet. 123: 779–791.

Camacho, C., G. Coulouris, V. Avagyan, N. Ma, J. Papadopoulos et al., 2009 BLAST+: Architecture and applications. BMC Bioinformatics 10:.

Cavallini, A., L. Natali, A. Zuccolo, T. Giordani, I. Jurman et al., 2010 Analysis of transposons and repeat composition of the sunflower (Helianthus annuus L.) genome. Theor. Appl. Genet. 120: 491–508.

Chen, Z. J., 2007 Genetic and epigenetic mechanisms for gene expression and phenotypic variation in plant polyploids. Annu. Rev. Plant Biol. 58: 377–406.

De Coster, W., S. D’Hert, D. T. Schultz, M. Cruts, and C. Van Broeckhoven, 2018 NanoPack: Visualizing and processing long-read sequencing data. Bioinformatics 34: 2666–2669.

Dai, S.-L., W.-K. Wang, M.-X. Li, and Y.-X. Xu, 2005 Phylogenetic Relationship of Dendranthema (DC.) Des Moul. Revealed by Fluorescent In Situ Hybridization. J. Integr. Plant Biol. 47: 783–791.

van Dijk, E. L., Y. Jaszczyszyn, D. Naquin, and C. Thermes, 2018 The Third Revolution in Sequencing Technology. Trends Genet. 34: 666–681.

Dowrick, G. J., 1952 The chromosomes of Chrysanthemum, I: the species. Heredity (Edinb). 6: 365–375.

Edger, P. P., T. J. Poorten, R. VanBuren, M. A. Hardigan, M. Colle et al., 2019 Origin and evolution of the octoploid strawberry genome. Nat. Genet. 51: 541–547.

Flynn, J. M., R. Hubley, C. Goubert, J. Rosen, A. G. Clark et al., 2020 RepeatModeler2 for automated genomic discovery of transposable element families. Proc. Natl. Acad. Sci. U. S. A. 117: 9451–9457.

Fonseca, N., and J. Manning fastq_utils. GitHub.

Garnatje, T., M. Á. Canela, S. Garcia, O. Hidalgo, J. Pellicer et al., 2011 GSAD: A genome size in the Asteraceae database. Cytom. Part A 79 A: 401–404.

van Geest, G., P. M. Bourke, R. E. Voorrips, A. Marasek-Ciolakowska, Y. Liao et al., 2017a An ultra-dense integrated linkage map for hexaploid chrysanthemum enables multi-allelic QTL analysis. Theor. Appl. Genet. 130: 2527–2541.

van Geest, G., R. E. Voorrips, D. Esselink, A. Post, R. G. F. Visser et al., 2017b Conclusive evidence for hexasomic inheritance in chrysanthemum based on analysis of a 183 k SNP array. BMC Genomics 18: 1–12.

Giordani, T., A. Cavallini, and L. Natali, 2014 The repetitive component of the sunflower genome. Curr. Plant Biol. 1: 45–54.

Götz, S., J. M. García-Gómez, J. Terol, T. D. Williams, S. H. Nagaraj et al., 2008 High-throughput functional annotation and data mining with the Blast2GO suite. Nucleic Acids Res. 36: 3420–3435.

Gurevich, A., V. Saveliev, N. Vyahhi, and G. Tesler, 2013 QUAST: Quality assessment tool for genome assemblies. Bioinformatics 29: 1072–1075.

Haas, B. J., S. L. Salzberg, W. Zhu, M. Pertea, J. E. Allen et al., 2008 Automated eukaryotic gene structure annotation using EVidenceModeler and the Program to Assemble Spliced Alignments. Genome Biol. 9: R7.

Hirakawa, H., K. Sumitomo, T. Hisamatsu, S. Nagano, K. Shirasawa et al., 2019 De novo whole-genome assembly in Chrysanthemum seticuspe, a model species of Chrysanthemums, and its application to genetic and gene discovery analysis. DNA Res. 26: 195–203.

Jones, P., D. Binns, H.-Y. Chang, M. Fraser, W. Li et al., 2014 InterProScan 5: genome-scale protein function classification. Bioinformatics 30: 1236–1240.

Kim, D., L. Song, F. P. Breitwieser, and S. L. Salzberg, 2016 Centrifuge: rapid and accurate classificaton of metagenomic sequences, version 1.0.4_beta. bioRxiv 26: 054965.

Klein, S. J., and R. J. O’Neill, 2018 Transposable elements: genome innovation, chromosome diversity, and centromere conflict. Chromosom. Res. 26: 5–23.

Kyriakidou, M., H. H. Tai, N. L. Anglin, D. Ellis, and M. V. Strömvik, 2018 Current strategies of polyploid plant genome sequence assembly. Front. Plant Sci. 871: 1660.

Li, H., 2013 Aligning sequence reads, clone sequences and assembly contigs with BWA-MEM.

Li, H., 2018 Minimap2: Pairwise alignment for nucleotide sequences. Bioinformatics 34: 3094–3100.

Li, F., G. Fan, C. Lu, G. Xiao, C. Zou et al., 2015 Genome sequence of cultivated Upland cotton (Gossypium hirsutum TM-1) provides insights into genome evolution. Nat. Biotechnol. 33: 524–530.

Liu, Y., 2020 Genetic structure and phenotypic differences among and within extant populations of Chrysanthemum arcticum L. and C. a. subsp. arcticum: University of Minnesota.

Liu, P. L., Q. Wan, Y. P. Guo, J. Yang, and G. Y. Rao, 2012 Phylogeny of the Genus Chrysanthemum L.: Evidence from Single-Copy Nuclear Gene and Chloroplast DNA Sequences. PLoS One 7:.

Liu, H., S. Wu, A. Li, and J. Ruan, 2021 SMARTdenovo: a de novo assembler using long noisy reads. Gigabyte 2021: 1–9.

Liu, B., J. Yan, W. Li, L. Yin, P. Li et al., 2020 Mikania micrantha genome provides insights into the molecular mechanism of rapid growth. Nat. Commun. 11: 340.

Lukaszewski, A. J., A. Alberti, A. Sharpe, A. Kilian, A. M. Stanca et al., 2014 A chromosome-based draft sequence of the hexaploid bread wheat (Triticum aestivum) genome. Science (80-.). 345:.

Ma, Y. P., M. M. Chen, J. X. Wei, L. Zhao, P. L. Liu et al., 2016 Origin of Chrysanthemum cultivars - Evidence from nuclear low-copy LFY gene sequences. Biochem. Syst. Ecol. 65: 129–136.

Marçais, G., and C. Kingsford, 2011 A fast, lock-free approach for efficient parallel counting of occurrences of k-mers. Bioinformatics 27: 764–770.

Natali, L., R. M. Cossu, E. Barghini, T. Giordani, M. Buti et al., 2013 The repetitive component of the sunflower genome as shown by different procedures for assembling next generation sequencing reads. BMC Genomics 14: 686.

Nguyen, T. K., S. T. T. Ha, and J. H. Lim, 2020 Analysis of chrysanthemum genetic diversity by genotyping-by-sequencing. Hortic. Environ. Biotechnol. 61: 903–913.

Okonechnikov, K., A. Conesa, and F. García-Alcalde, 2016 Qualimap 2: Advanced multi-sample quality control for high-throughput sequencing data. Bioinformatics 32: 292–294.

Ou, S., J. Chen, and N. Jiang, 2018 Assessing genome assembly quality using the LTR Assembly Index (LAI). Nucleic Acids Res. 46: e126.

Palmer, J. M., 2017 Funannotate: Eukaryotic Genome Annotation Pipeline.

Peng, Y., Z. Lai, T. Lane, M. Nageswara-Rao, M. Okada et al., 2014 De Novo Genome Assembly of the Economically Important Weed Horseweed Using Integrated Data from Multiple Sequencing Platforms. PLANT Physiol. 166: 1241–1254.

Pflug, J. M., V. R. Holmes, C. Burrus, J. Spencer Johnston, and D. R. Maddison, 2020 Measuring genome sizes using read-depth, k-mers, and flow cytometry: Methodological comparisons in beetles (Coleoptera). G3 Genes, Genomes, Genet. 10: 3047–3060.

Ramírez-Sánchez, O., P. Pérez-Rodríguez, L. Delaye, and A. Tiessen, 2016 Plant Proteins Are Smaller Because They Are Encoded by Fewer Exons than Animal Proteins. Genomics, Proteomics Bioinforma. 14: 357–370.

Roach, M. J., S. A. Schmidt, and A. R. Borneman, 2018 Purge Haplotigs: Allelic contig reassignment for third-gen diploid genome assemblies. BMC Bioinformatics 19: 460.

Ruan, J., H. Li, D. Li, and W. Lin SMARTdenovo. GitHub.

Scaglione, D., S. Reyes-Chin-Wo, A. Acquadro, L. Froenicke, E. Portis et al., 2016 The genome sequence of the outbreeding globe artichoke constructed de novo incorporating a phase-aware low-pass sequencing strategy of F 1 progeny. Sci. Rep. 6: 1–17.

Schnable, P. S., D. Ware, R. S. Fulton, J. C. Stein, F. Wei et al., 2009 The B73 maize genome: Complexity, diversity, and dynamics. Science (80-.). 326: 1112–1115.

Shen, Q., L. Zhang, Z. Liao, S. Wang, T. Yan et al., 2018 The Genome of Artemisia annua Provides Insight into the Evolution of Asteraceae Family and Artemisinin Biosynthesis. Mol. Plant 11: 776–788.

Simão, F. A., R. M. Waterhouse, P. Ioannidis, E. V. Kriventseva, and E. M. Zdobnov, 2015 BUSCO: Assessing genome assembly and annotation completeness with single-copy orthologs. Bioinformatics 31: 3210–3212.

Song, C., Y. Liu, A. Song, G. Dong, H. Zhao et al., 2018 The Chrysanthemum nankingense Genome Provides Insights into the Evolution and Diversification of Chrysanthemum Flowers and Medicinal Traits. Mol. Plant 11: 1482–1491.

Tanaka, R., 1960 On the Speciation and Karyotypes in Diploid and Tetraploid Species of Chrysanthemum. Cytologia (Tokyo). 25: 43–58.

Tanaka, R., and N. Shimotomai, 1968 A Cytogenetic Study on the F1 Hybrid of Chrysanthemum makinoi×Ch. vulgare. Cytologia (Tokyo). 33: 241–245.

Tang, H., X. Zhang, C. Miao, J. Zhang, R. Ming et al., 2015 ALLMAPS: Robust scaffold ordering based on multiple maps. Genome Biol. 16: 3.

Todorovska, E., 2007 Retrotransposons and their role in plant—genome evolution. Biotechnol. Biotechnol. Equip. 21: 294–305.

Vurture, G. W., F. J. Sedlazeck, M. Nattestad, C. J. Underwood, H. Fang et al., 2017 GenomeScope: Fast reference-free genome profiling from short reads. Bioinformatics 33: 2202–2204.

Wang, H., X. Qi, R. Gao, J. Wang, B. Dong et al., 2014 Microsatellite polymorphism among Chrysanthemum sp. polyploids: The influence of whole genome duplication. Sci. Rep. 4: 1–8.

Warren, R. L., L. Coombe, H. Mohamadi, J. Zhang, B. Jaquish et al., 2019 ntEdit: scalable genome sequence polishing. Bioinformatics 35: 4430–4432.

Wick, R. R., 2019 Filtlong: quality filtering tool for long reads.

Wick, R. R., 2018 Porechop: adapter trimmer for Oxford Nanopore reads.

Wick, R. R., L. M. Judd, C. L. Gorrie, and K. E. Holt, 2017 Unicycler: Resolving bacterial genome assemblies from short and long sequencing reads (A. M. Phillippy, Ed.). PLOS Comput. Biol. 13: e1005595.

Won, S. Y., Y. J. Hwang, J. A. Jung, J. S. Kim, S. H. Kang et al., 2018a Identification of repetitive DNA sequences in the Chrysanthemum boreale genome. Sci. Hortic. (Amsterdam). 236: 238–243.

Won, S. Y., J.-A. Jung, and J. S. Kim, 2018b The complete chloroplast genome of Chrysanthemum boreale (Asteraceae). Mitochondrial DNA Part B 3: 549–550.

Won, S. Y., J.-A. Jung, and J. S. Kim, 2018c The complete mitochondrial genome sequence of Chrysanthemum boreale (Asteraceae). Mitochondrial DNA Part B 3: 529–530.

Zhanga, H., B. Zhu, B. Qi, X. Gou, Y. Dong et al., 2014 Evolution of the BBAA component of bread wheat during its history at the allohexaploid level. Plant Cell 26: 2761–2776.

